# Lithium Chloride Inhibits Iron Dysregulation and Ferroptosis in Induced Pluripotent Stem Cells with ApoE4/E4 from a sporadic Alzheimer’s disease patient

**DOI:** 10.1101/2025.08.28.672956

**Authors:** Ying Wang, Samuel Anchipolovsky, Piplu Bhuiyan, Luna Sato, Ge Liang, De-Maw Chuang, Huafeng Wei

## Abstract

Alzheimer’s disease (AD), particularly its sporadic form (SAD, 95% AD patients), is strongly associated with the apolipoprotein E4 ApoE4 genotype and characterized by oxidative stress, iron dysregulation, and increased susceptibility to ferroptosis. Lithium, a well-established neuroprotective agent, has shown potential to mitigate several pathological mechanisms in AD, including ferroptosis. This study investigates the therapeutic potential of lithium chloride in human induced pluripotent stem cells (iPSCs) derived from a SAD patient with ApoE4/E4 genotype, and compared effects with those of isogenic gene-edited ApoE3/E3 control. Lithium treatment significantly improved cell viability in ApoE4/E4 iPSCs. It also reversed key ferroptosis phenotypes, including elevated cytosolic Fe^2+^, increased expression of divalent metal transporter 1, reduced level of glutathione peroxidase 4, enhanced lipid peroxidation, and excessive ROS production. Moreover, lithium normalized mitochondrial respiration and reduced proton leak, indicating preservation of mitochondrial function and protection against mitochondrial damage and cell death. Lithium also reduced the expression of type 1 InsP3 receptor (InsP3R-1) protein, a Ca^2+^ channel located on the endoplasmic reticulum (ER) membrane. Together, these findings highlight lithium’s inhibition of ferroptosis through modulation of iron metabolism, antioxidant defenses, and inhibition of disrupted Ca^2+^ signaling. Given its demonstrated efficacy in reversing ApoE4-driven cellular vulnerabilities, lithium salt warrants further investigation for the treatment of AD.

## Introduction

Alzheimer’s disease (AD) is a progressive neurodegenerative disorder characterized by cognitive decline, neuronal loss and the accumulation of amyloid-beta plaques together with hyperphosphorylated tau. Sporadic AD (SAD), the most common form, is strongly influenced by the apolipoprotein E4 (ApoE4) allele, which exacerbates the AD pathology through multiple pathways including oxidative stress, mitochondrial dysfunction, iron dysregulation, and Ca^2+^ homeostasis disruption^1,2^. Ferroptosis, an iron-dependent form of programmed cell death, has emerged as a critical contributor to AD-associated neurodegeneration and associated cognitive dysfunction^3^. This process is driven by elevated intracellular iron (Fe^2+^) levels, mediated by upregulation of divalent metal transporter 1 (DMT1) expression, resulting in reactive oxygen species (ROS) accumulation, lipid peroxidation, and cell death^4,5^. ApoE4-expressing cells exhibit heightened susceptibility to ferroptosis due to reduced glutathione peroxidase 4 (GPX4) activity, a key enzyme mitigating lipid peroxidation^6,7^. ApoE4 has been demonstrated to exacerbate ferroptosis in AD^6,7^. Moreover, both Ca^2+^ and Fe^2+^ dysregulation contribute to ferroptosis in AD^8,9^.

Lithium, a well-established mood stabilizer, exhibits neuroprotective effects in preclinical models of neurodegenerative diseases, notably AD, via multiple mechanisms including modulating oxidative stress, excitotoxicity, Ca^2+^ dysregulation, and cellular signaling pathways^10-12^. A recent study indicates that a deficiency of lithium in the brain increases the risk of developing AD^13^. It is generally believed that inhibition of glycogen synthase kinase-3 and inositol monophosphatase are two primary initial events by which lithium exerts neuroprotection^14^. Lithium enhances GPX4 expression and activity through inhibition of glycogen synthase kinase-3 and activation of the nuclear factor erythroid 2-related factor 2 pathway, bolstering antioxidant defenses^15-17^. Conversely, inhibition of inositol monophosphatase by lithium triggers the induction of autophagy, a process prominently involved in the removal of pathological proteins and frequently dysregulated in brain disorders including AD^14,18^. Additionally, lithium reduces iron dysregulation by downregulating DMT1 expression, thereby limiting Fe^2+^ accumulation and ferroptosis^5^. Furthermore, lithium ameliorates ApoE4- and N-methyl-D-asparte (NMDA) receptor-mediated Ca^2+^ dysregulation by inhibiting NMDA receptor-mediated Ca^2+^ influx and Src tyrosine kinase activity, mitigating excitotoxicity and oxidative stress that exacerbates ferroptosis in AD^19,20^. Lithium provides synergistic inhibition of ferroptosis via modulation of tau phosphorylation and MAPK signaling in a cell model of AD^21^. On the other hand, lithium enhances ferroptosis in melanoma cells^22^. Induced pluripotent stem cells (iPSCs) derived from SAD patients with ApoE4/E4 genotype provide a robust model to study these mechanisms, as they recapitulate disease-specific phenotypes, including elevated Fe^2+^, ROS, mitochondrial dysfunction, and Ca^2+^ dysregulation^23,24^.

This study investigates lithium’s role in inhibiting ferroptosis in iPSCs derived from a SAD patient with parental ApoE4/E4 genotype, in comparison to the gene-edited isogenic ApoE3/E3. This is an excellent translational cell model of SAD to test a new drug’s therapeutic efficacy and underlying mechanisms, considering that ApoE4 is one of the primary risk factors for SAD. We hypothesize that lithium exerts cytoprotective effects in these iPSCs, at least in part, by inhibition of ferroptosis. In support of this hypothesis, we show here that lithium reduces DMT1-mediated Fe^2+^accumulation and ameliorates mitochondrial dysfunction and other detrimental cellular events to prevent cell death. These findings provide insight into lithium’s therapeutic potential as a cytoprotective agent against ferroptosis in AD.

## Materials and Methods

### Cell Culture

Cell models featuring the primary susceptibility factor for sporadic Alzheimer’s disease (SAD), namely ApoE4, and induced pluripotent stem cells (iPSCs) were procured from the laboratory of Dr. Li-Huei Tsai^1^, Massachusetts Institute of Technology. ApoE3/E3 cells were derived from CRISPR/Cas9 editing of SAD iPSCs homozygous for ApoE4/E4 (AG10788 cell line from an 87-year-old familial female AD patient). The human iPSCs were cultivated on Matrigel-coated plates from BD Biosciences, USA, utilizing mTeSR™ Plus medium (catalog No. 100-0276, Stem Cell Technologies, Canada), and were incubated in a 5% CO2 humidified atmosphere at 37°C. The culture medium underwent daily replacement. Before experiments, a regular assessment of cell health was conducted, and any unhealthy cells, such as nonadherent cells, were systematically eliminated. The cells were randomly allocated into two distinct experimental groups.

### Cell viability assessment

To evaluate cell viability in distinct wells of 96-well plates, we employed the 3-(4,5-dimethylthiazol-2-yl)-2,5-diphenyltetrazolium bromide (MTT; Sigma–Aldrich, USA) reduction assay at a 24-hour time point, using a previously established protocol25,26. Following a wash with phosphate-buffered saline, the samples underwent incubation with a fresh culture medium containing MTT (0.5 mg/ml in the medium) at 37°C for 4 hours in the absence of light. Subsequently, the medium was aspirated, and formazan reaction products were solubilized with dimethyl sulfoxide. The resulting absorbance was measured at 570 nm using a plate reader (Synergy H1 microplate reader, BioTek, CA, USA).

### Measurement of mitochondrial function by using a Seahorse XFp Extracellular Flux Analyzer

In the pursuit of measuring oxygen consumption rate (OCR) and proton leak in iPSCs, a sophisticated methodology was employed, utilizing Seahorse XFp miniplates, the cell energy phenotype test and the Seahorse XFp Analyzer software. iPSCs were meticulously cultured and plated in Seahorse XFp miniplates to ensure optimal cell adherence under conditions that sustain cellular health. After cell adherence, the Seahorse XFp Analyzer was utilized to conduct the cell energy phenotype test, a dynamic assay providing real-time insights into critical parameters such as OCR and proton leak, pivotal indicators of cellular bioenergetics. The Seahorse XFp Analyzer diligently monitors iPSC OCR, reflecting the rate at which cells consume oxygen during oxidative phosphorylation. Measurements were conducted under basal conditions and the reactions were augmented by the sequential addition of mitochondrial modulators, including oligomycin, carbonyl cyanide-4 (trifluoromethoxy) phenylhydrazone (FCCP) and antimycin A/rotenone. These additions facilitated the assessment of key mitochondrial functions, encompassing ATP production, maximal respiration and spare respiratory capacity. Proton leak, a pivotal facet of mitochondrial function, underwent specific evaluation, denoting the passive leakage of protons across the inner mitochondrial membrane, contributing to basal OCR. This provided insight into mitochondrial uncoupling and energy inefficiencies. The Seahorse XFp Analyzer, coupled with appropriate inhibitors, facilitated the precise quantification of proton leak in iPSCs. Data analysis was performed via web-based Seahorse software and results were normalized based on cell viability upon completion of assay.

### Intracellular MDA concentration assay

Lipid peroxidation levels were evaluated by quantifying the cellular content of malondialdehyde (MDA) using the colorimetric Lipid Peroxidation Assay kit (ab118970, Abcam, Cambridge, UK), following the manufacturer’s instructions. In this procedural framework, MDA within the sample underwent a reaction with thiobarbituric acid (TBA), producing an MDA-TBA adduct, the colorimetric quantification of which was subsequently expressed as MDA levels. Specifically, following a 24-hour drug treatment, iPSCs were collected and homogenized in 303 µL of MDA lysis solution. To generate the MDA-thiobarbituric acid (TBA) adduct, 600 µL of TBA reagent was added to 200 µL of the sample. The resultant mixture was incubated at 95 °C for 60 minutes and then cooled on ice. The absorbance of the resulting supernatant, containing the MDA-TBA adduct, was measured at 532 nm using a microplate reader, and MDA levels (µM) were calculated by comparing the absorbance of each sample to a standard.

### Intracellular ferrous iron concentration assay

The Cell Ferrous Iron Colorimetric Assay Kit (EBC-K881-M, Elabscience, Houston, Texas, USA) was employed for the quantification of intracellular ferrous iron levels. Approximately 1 × 10^6^ cells were extracted and homogenized on ice for 10 minutes with 200 µL of lysis buffer, followed by centrifugation at 15,000× g for 10 minutes to collect the supernatant. Subsequently, 80 µL of the supernatant underwent treatment with the iron probe or the control reagent for 10 minutes at 37°C. A Multiscan FC plate reader (Synergy H1 microplate reader, BioTek, USA) was utilized to measure the absorbance at 593 nm. The relative ferrous iron levels were determined as the difference between the ferrous iron contents of the experimental and control groups. A standard curve for cell ferrous iron was employed to calculate the cellular ferrous iron content.

### Intracellular ROS assay

To assess intracellular levels of reactive oxygen species (ROS), we conducted the 2′,7′-Dichlorofluorescein diacetate, Diacetyldichlorofluorescein (DCFDA) assay following a meticulous microplate protocol. Initially, iPSCs were seeded and allowed to be attached in a 96-well plate. Following this, a washing step with an appropriate buffer was implemented to eliminate extraneous substances that could potentially interfere with the assay. Subsequently, the iPSCs were subjected to staining with DCFDA, a fluorogenic dye sensitive to ROS. The staining duration was 45 minutes to ensure optimal dye penetration. Post-staining, thorough washing was performed to remove excess, unincorporated dye, ensuring a specific and accurate assessment of intracellular ROS levels. The stained iPSC samples remained in the 96-well plate, distributed evenly across the wells. Employing a plate reader (Synergy H1 microplate reader, BioTek, USA), fluorescence measurements were then taken at the specified excitation and emission wavelengths (Ex/Em = 485/535 nm) for DCFDA. The recorded fluorescence readings for each well were subsequently analyzed, providing a quantitative measure of intracellular ROS levels in the iPSC population.

### Western blotting analysis

The Western blotting procedure adhered to established standards. Total protein extracts from iPSCs were acquired by lysing the cell in cold RIPA buffer (#9806S, Cell Technology, CA, USA) supplemented with protease inhibitor cocktails (P8340 Roche). Following centrifugation, the supernatant was collected, and total protein quantification was performed using a bicinchoninic acid protein assay kit (Thermo Scientific, MA, USA). Equal quantities of protein for each lane were loaded and subjected to separation on a 4-20% TGX gel (BioRad, CA, U.S.A.) through electrophoresis. Post-electrophoresis, the proteins were transferred onto a polyvinylidene fluoride membrane. Subsequently, the membranes were obstructed with 5% bovine serum albumin dissolved in phosphate-buffered saline-T for 1 hour at room temperature and then probed with the primary antibody (GPX4, DMT-1 (20507-1-AP, Dilution, 1:2000), InsP_3_R-1 (Catalog # PA1-901, Invitrogen, CA, U.S.A, Dilution, 1:1000) at 4°C overnight. After thorough washing with phosphate-buffered saline-T, the membranes underwent incubation with secondary antibodies (horseradish peroxidase– conjugated anti-rabbit and anti-mouse IgG) at 1:10,000 dilutions, with GAPDH serving as a loading control. Signals were detected through an enhanced chemiluminescence detection system (Millipore, MA, USA) and quantified using scanning densitometry.

## Data analysis and statistics

Prior to the study, no formal statistical power analysis was conducted, and the determination of the sample size was based on our previous experience with a similar experimental design. The normality of data distribution was evaluated using the Tukey test and Fisher test, which guided the decision on whether to employ parametric or nonparametric statistical analyses. Variables meeting the assumptions for parametric analysis were presented as mean ± SD and subjected to either one-way or two-way analysis of variance, followed by Sidak’s post hoc analysis. Consideration was given to factors such as the nature of factors (e.g., repeated measures) and the grouping of these factors in the analysis of variance. Variables meeting the assumptions for nonparametric analysis were scrutinized using the Tukey multiple comparisons test and Fisher’s Exact test. Statistical analyses and graphical representations were conducted using GraphPad Prism software (GraphPad Software, Inc., MA, USA). A significance threshold of P < 0.05 was employed to determine statistical significance.

## Results

### Lithium enhances viability in ApoE4/E4 iPSC-derived cells by restoring intracellular Ca^2+^ homeostasis

To evaluate the cytoprotective potential of lithium in SAD-relevant cell lines, cell viability was assessed in parental ApoE4/E4 and isogenic gene-modified ApoE3/E3 iPSCs using the MTT assay, which measures mitochondrial dehydrogenase activity. ApoE4/E4 cells demonstrated significantly reduced viability in comparison to ApoE3/E3 controls, consistent with genotype-associated cellular vulnerability (Fig. 1A). Treatment with 0.25 or 1.25 mM lithium chloride for 24 hours resulted in a marked improvement in ApoE4/E4 cell viability (Fig. 1A). Thus, lithium confers protective effects against ApoE4-mediated cytotoxicity. To determine if elevation of cytosolic Ca^2+^ concentration plays a role in ApoE4 mediated iPSCs damage, we used BAPTA-AM to ameliorate the increase of cytosolic Ca^2+^, which dose-dependently promoted ApoE4/E4 iPSC survival (Fig. 1B). On the other hand, NMDA, which selectively activates the NMDA subtype of glutamate receptors on plasma membrane and increases cytosolic Ca^2+^ concentration, induced more potent and robust cell damages in ApoE4/E4 than in ApoE3/E3 iPSCs (Fig. 1C), and these NMDA-induced cytotoxic effects were abolished by lithium treatment (Fig. 1D).

**Figure 1.**
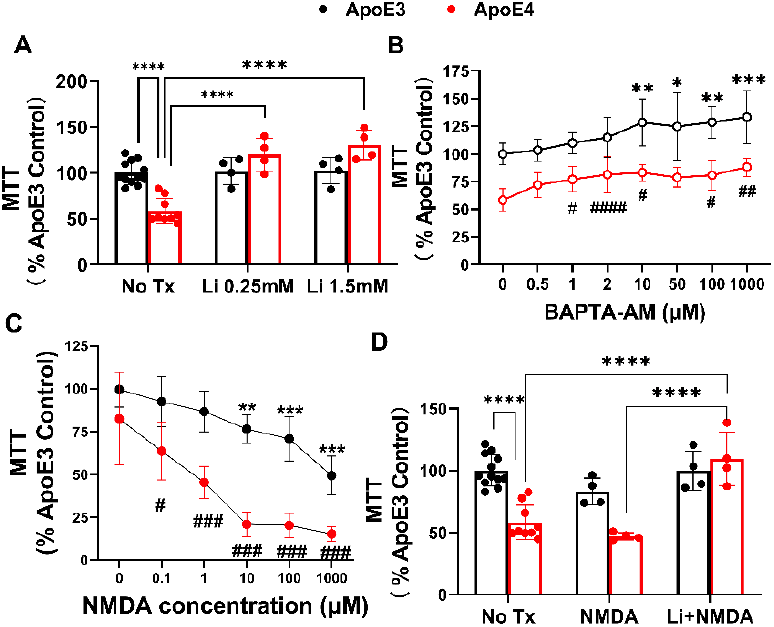
Treatment with lithium chloride inhibits ApoE4/NMDA-mediated mitochondrial and cell damage. iPSCs with parental ApoE4/E4 (ApoE4) from a SAD patient or its isogenic gene-edited ApoE3/E3 (ApoE3) were treated with 0.25 or 1.5 mM lithium chloride (Li) **(A)** or intracellular Ca^2+^ chelator to inhibit elevation of cytosolic Ca^2+^ concentration (BAPTA-AM) (**B)** or N-methyl-D-aspartate (NMDA) (**C**) at different concentrations for 24 hours. (**D)** Cells were pretreated with Li (0.25 mM) for one hour and then co-treated with NMDA (1 μM) for 24 hours. Lower MTT corresponds to greater levels of mitochondrial and cell damage. Means ± SD from 6 repeats of 4 to 8 separate experiments (N= 24-32 repeats). Two-way ANOVA followed by Tukey Multiple comparison tests (MCT). **** P<0.0001 (**A, D)**. *, **, ***, **** P<0.05, P<0.01, P<0.001 and P<0.0001 compared to control in same type of cells respectively (**B, C**). ^#, ##, ###, ####^, P<0.05, P<0.01, P<0.001 and P<0.0001 compared to ApoE3 control cells respectively **(B, C)**. Tx: Treatment **(A, D)**.

### Lithium suppresses ApoE4/NMDA-mediated increases of type 1 InsP_3_R proteins

To determine whether lithium modulates Ca^2+^signaling through regulation of type 1 inositol 1,4,5-trisphosphate receptor (InsP3R-1) expression, protein levels of InsP_3_R-1 were assessed using Western blotting (Fig. 2). ApoE4/E4 cells exhibited a three to four-fold increase in InsP_3_R-1 protein levels compared to ApoE3/E3 controls. Lithium treatment completely blocked the InsP_3_R-1 protein upregulation in ApoE4/E4 cells, supporting that lithium suppresses ApoE4-associated upregulation of this intracellular Ca^2+^ channel. This finding suggests that lithium inhibits InsP_3_R-1 expression driven by ApoE4 genotype. This result highlights a potential mechanism by which lithium mitigates Ca^2+^ dysregulation in ApoE4-mediated cell damage.

**Figure 2.**
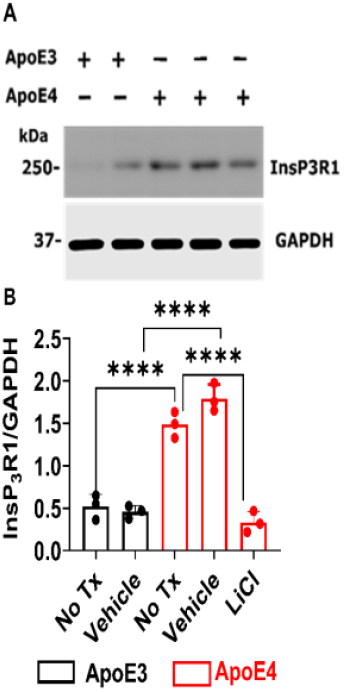
Lithium blocks ApoE4-mediated robust elevation of type 1 InsP_3_R (InsP_3_R1) proteins. iPSCs with parental ApoE4/E4 (ApoE4) from a SAD patient or its isogenic gene-edited ApoE3/E3 (ApoE3) were treated with lithium chloride (Li, 0.25 mM) or vehicle (H_2_O) or without any treatment (TX) for 24 hours. Western Blot. Means±SD, N=3 separate experiments. GAPDH protein level was used as a loading control. Two-way ANOVA followed by Tukey’s multiple comparison test. ****P<0.0001.

### Lithium reduces intracellular Fe^2+^ accumulation in ApoE4/E4 cells

To investigate lithium’s influence on iron homeostasis, a major facilitator of cell death by ferroptosis, cytosolic ferrous iron (Fe^2+^) levels were measured using a colorimetric iron assay kit. ApoE4/E4 cells exhibited significantly elevated cytosolic Fe^2+^ concentrations relative to ApoE3/E3 controls (Fig. 3A). Pretreatment with lithium normalized Fe^2+^ levels in ApoE4/E4 cells, restoring them to near-control levels (Fig. 3A). Furthermore, lithium treatment completely suppressed the additional increase in cytosolic Fe^2+^ concentration induced by exposure to NMDA (Fig. 3B) and RSL3, a GPX4 inhibitor and ferroptosis inducer (Fig. 3C), in ApoE4/E 4iPSCs.

**Figure 3.**
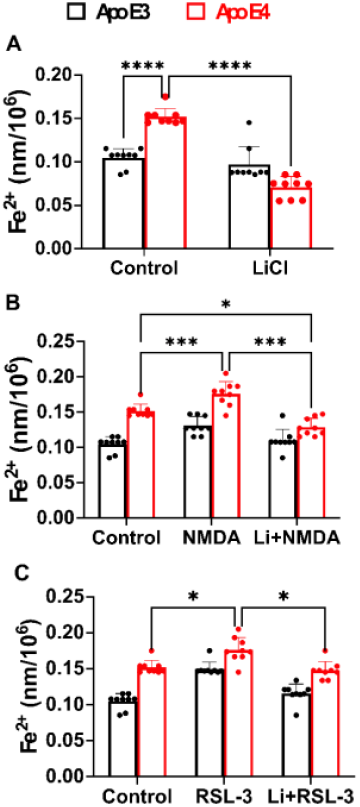
Lithium completely suppresses ApoE4/NMDA/RSL-3 mediated increases of cytosolic Fe^2+^ concentrations. iPSCs with parental ApoE4/E4 (ApoE4) from a SAD patient or its isogenic gene-edited ApoE3/E3 (ApoE3) were treated with lithium chloride (Li, 0.25 mM) (**A, B and C)**, with or without 1 µM NMDA (**B**) or 5 μM RSL3 (**C)** for 24 hours. The data is presented as the mean ± SD from three independent experiments with a total of 9 repeats (N=9) (**A, B, C**). Statistical analysis using two-way analysis of variance followed by Tukey multiple comparison tests (MCT). *P<0.05, ***P <0.001, ****P < 0.0001. NMDA: The N-methyl-D-aspartate, RSL3, Ras, selective lethal 3, a selective inhibitor of GPX4.

### Lithium blocks DMT1 overexpression in ApoE4/E4 cells

To determine the potential mechanism through which lithium affects cytosolic iron concentration, we measured the changes in protein level of the iron transporter divalent Metal Transporter 1 (DMT1). ApoE4/E4 cells exhibited a pronounced upregulation of DMT1 compared to ApoE3/E3 controls (Fig. 4). Lithium treatment markedly reduced the DMT1 protein upregulation in ApoE4/E4 cells to the level in ApoE3/E3 iPSCs.

**Figure 4.**
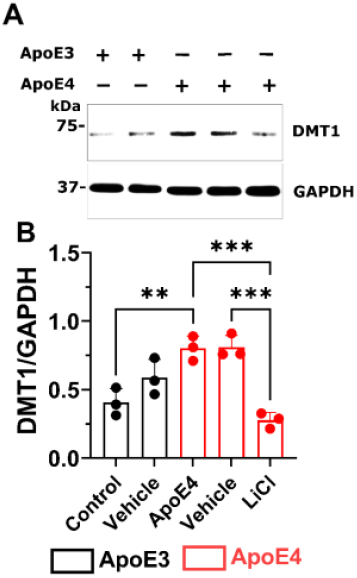
Lithium abolishes ApoE4-mediated pathological elevation of divalent metal transporter 1 (DMT1) proteins. iPSCs with parental ApoE4/E4 (ApoE4) from a SAD patient or its isogenic gene-edited ApoE3/E3 (ApoE3) were treated with lithium chloride (Li, 0.25 mM) or vehicle (H_2_O), or without any treatment (No Tx) for 24 hours. Western blot with primary antibody against DMT1. Means±SD. Two-way ANOVA followed by multiple Tukey test. ***P<0.001.

### Lithium restores antioxidant defense and reduces oxidative stress

Glutathione Peroxidase 4 (GPX4), an antioxidant enzyme protecting cells from oxidative stress, is typically downregulated in ferroptosis. GPX4 expression was significantly reduced in ApoE4/E4 cells compared to ApoE3/E3 controls (Fig. 5). Lithium treatment restored GPX4 expression to levels comparable to those in ApoE3/E3 cells (Fig. 5A), indicating a reconstitution of lipid peroxide repair capacity.

**Figure 5.**
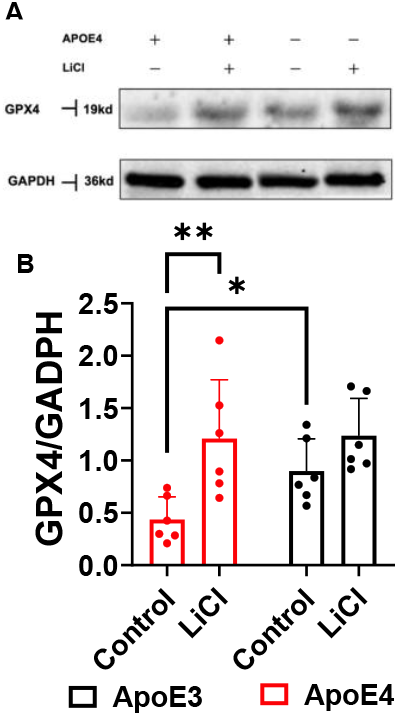
Lithium treatment restores ApoE4-mediated pathological decrease of glutathione peroxidase 4 protein (GPX4) expression. iPSCs with parental ApoE4/E4 (ApoE4) from a SAD patient or its isogenic gene-edited ApoE3/E3 (ApoE3) were treated with lithium chloride (Li, 0.25 mM) for 24 hours, then cell proteins collected to determine GPX4 protein, an enzyme that removes reactive oxygen species (ROS). **A**. Representative of Western blotting images of GPX4 proteins expression under different experimental conditions in two types of cells. (**B)** Statistical analysis for Western blot data. Data is presented as mean ± SD, from 5-6 separate experiments (N=5 or 6). Two-way ANOVA followed by Tukey multiple comparison tests (MCT). *P < 0.05; **P < 0.01.

To determine intracellular oxidative stress level, we measured change in malondialdehyde (MAD), a metabolite of lipid peroxidation and reactive oxygen species (ROS). Intracellular MDA concentration in ApoE4/E4 iPSCs was significantly higher than that in ApoE3/E3 iPSCs. Treatment with lithium chloride at 0.25 mM for 24 hours restored the elevated intracellular MDA concentration in ApoE4/E4 iPSCs to the level in ApoE3/E3 iPSCs (Fig. 6A). Treatment with 1 μM NMDA further increased intracellular MDA concentration in ApoE4/E4 iPSCs, which was significantly inhibited by lithium treatment (Fig. 6B). The intracellular ROS levels in ApoE4/E4 iPSCs tended to be higher than those in ApoE3/E3 iPSCs. RSL-3, an inhibitor of GPX4 (Fig. 6C), and NMDA (Fig. 6D) significantly increased intracellular ROS level in ApoE4/E4 but not ApoE3/E3 iPSCs. Pretreatment with lithium for 24 hours effectively ameliorated the RSL-3 (Fig. 6C) or NMDA (Fig. 6D) -induced pathological elevation of intracellular R in two types of cells. **(B)** Statistical analysis for Western blot data. Data is presented as mean ± SD, from 5-6 separate experiments (N=5 or 6). Two-way ANOVA followed by Tukey multiple comparison tests (MCT). *P < 0.05; **P < 0.01.

**Figure 6.**
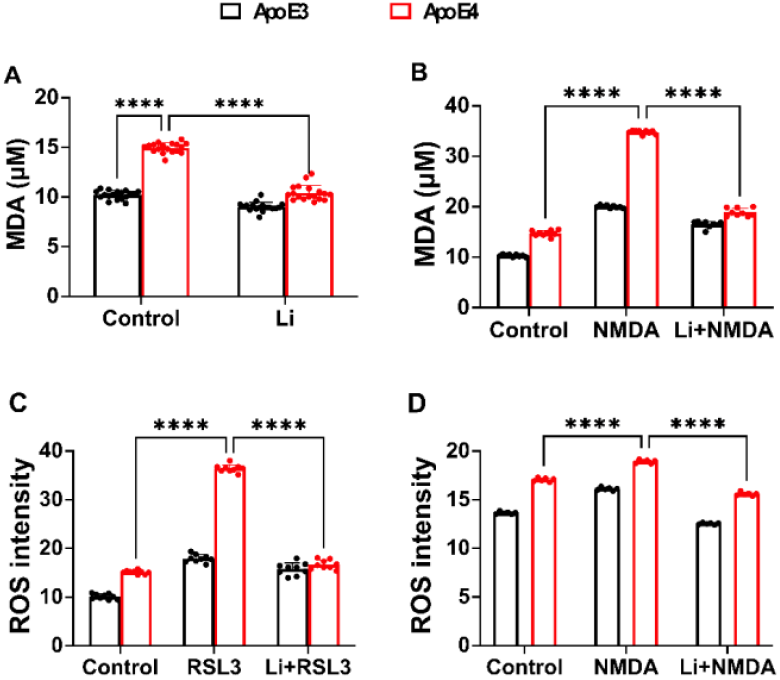
Lithium inhibits ApoE4/NMDA/RSL3-mediated elevation of intracellular reactive oxygen species (ROS) and lipid membrane oxidation metabolite malondialdehyde (MDA). iPSCs with parental ApoE4/E4 (ApoE4) from a SAD patient or its isogenic gene-edited ApoE3/E3 (ApoE3) were treated with lithium chloride (Li, 0.25 mM) for 24 hours (**A**), with or without co-treatments of NMDA (1 μM) (**B, D**) or RSL-3 (selective inhibitor of GPX4, **C**). Intracellular malondialdehyde (MDA), a by-product of lipid peroxidation, or reactive oxygen species (ROS) were then quantified. **(A, B)**. Reactive oxygen species (ROS) in cytosol were measured (**C, D**). Data represents Mean+SD from 16 (N=16, **A**), 12 (N=12, **B**), 16 (N=16, **C**) and 16 repeats (N=16, **D**) of 4 separate experiments. Two-way ANOVA followed by Tukey multiple comparison test. ^**\*\*\*\***^ P<0.0001 (**A, B, C, D**).

### Lithium normalizes mitochondrial oxygen consumption and reduces proton leak in ApoE4/E4 cells

To evaluate the impact of lithium on mitochondrial bioenergetics, oxygen consumption rate (OCR) and proton leak were measured using Seahorse metabolic flux analysis. ApoE4/E4 iPSCs exhibited elevated basal OCR, which was significantly exacerbated by RSL-3 (Fig. 7 A, B, C, E). Pretreatment with lithium chloride at 0.25 mM for 24 hours significantly inhibited RSL-3-induced elevation of basal OCR. Proton leak (an indication of mitochondrial and cell damage by apoptosis40) in ApoE4/E4 increased in comparison to ApoE3/E3 iPSCs, which was further significantly increased by RSL-3 (Fig. 7 A, B, D, F). Lithium chloride (0.25 mM) pretreatment for 24 hours abolished RSL-3-induced proton leaks in ApoE4/E4 iPSCs (Fig. 7D, F).

**Figure 7.**
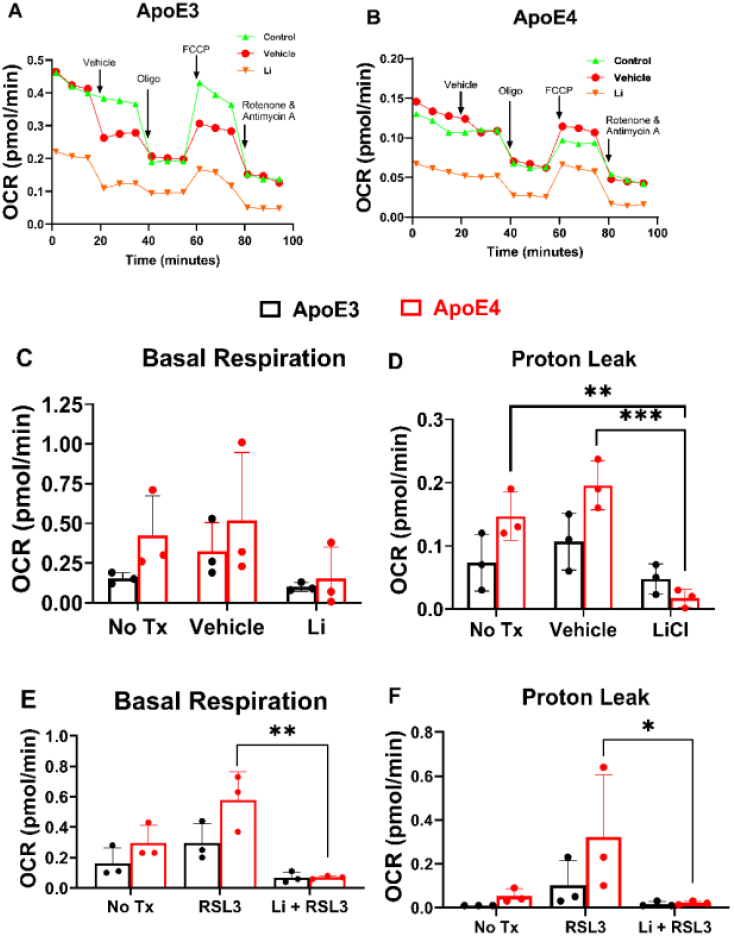
Lithium strongly inhibits ApoE4 or Ras-selective lethal 3 (RSL3)-mediated pathological elevation of basal mitochondrial oxygen consumption rate (OCR) and proton leak (apoptosis). iPSCs with parental ApoE4/E4 (ApoE4) from a SAD patient or its isogenic gene-edited ApoE3/E3 (ApoE3) were treated with lithium chloride (Li, 0.25 mM) for 24 hours, then challenged with 5 μM RSL3 or vehicle control for one hour. Seahorse XF Mito Stress assay was performed, with typical OCR tracing profiles depicted for ApoE3 **(A)** and ApoE4 **(B)**, and arrows indicating sequential addition of vehicle, oligomycin (Oligo), FCCP, and rotenone/antimycin. Basal respiration OCR was determined under treatment of LiCl or vehicle (**C**), with or without RSL-3 (**E**). Control cells received no treatment (No Tx). Mitochondrial proton leak as an apoptosis biomarker was calculated based on OCR in two types of cells under treatment of LiCl or vehicle (**D**), with or without treatment of RSL-3 (**F**). Control cells receive no treatment (No Tx). Data are presented as mean ± SD, N=3 separate experiments (**C-F**). Two-way ANOVA followed by Tukey multiple comparison test (MCT). *P < 0.05 (**F**); **P < 0.01 (**D, E**), ***P < 0.001 (**D**).

## Discussion

This study provides compelling evidence that lithium treatment protects ApoE4/E4 human iPSC-derived neurons from ferroptosis through multimodal mechanisms involving the regulation of iron, Ca^2+^ signaling and antioxidant defense.

Specifically, lithium blocked the upregulation of type 1 InsP_3_R proteins, reduced cytosolic Fe^2+^ accumulation, suppressed the increased iron transporter DMT1, restored GPX4 expression, normalized the mitochondrial function, and mitigated lipid peroxidation and ROS generation in ApoE4/E4 cells, with or without glutamate excitotoxicity and additional oxidative stress. These effects were accompanied by a significant increase in cell viability in the ApoE4/E4 cells, which is known to confer elevated susceptibility to oxidative and metabolic stressors^27-29^.

These findings expand upon a growing body of literature demonstrating lithium’s neuroprotective effects in models of neurodegeneration. Lithium has long been shown to exert neurotrophic and anti-apoptotic effects through inhibition of glycogen synthase kinase-3, enhancement of the signaling of brain-derived neurotrophic factor (BDNF) and other trophic factors, as well as modulation of mitochondrial function^14,30^. In the context of AD, lithium has been associated with reduced amyloid pathology, tau hyperphosphorylation, and neuroinflammation in both preclinical and clinical settings^21,31,32^. However, its role in regulating ferroptosis, a distinct form of regulated cell death driven by iron accumulation and lipid peroxidation, is only beginning to be elucidated^21^. Recent studies have implicated ferroptosis in the pathophysiology of AD, particularly in the presence of the ApoE4 genotype, which has been shown to disrupt lipid metabolism, increase oxidative stress, and promote iron accumulation^29,33,34^.

Our data are consistent with this genotype-dependent vulnerability and further demonstrate that lithium treatment can reverse these ferroptosis phenotypes in ApoE4/E4 iPSC. This suggests that lithium’s cytoprotection may extend beyond traditional anti-apoptotic pathways to include ferroptosis inhibition^21^.

Crucially, the present study identifies a novel mechanism by which lithium inhibits ferroptosis. Previous work has shown that excessive Ca^2+^ influx, such as that triggered by glutamatergic excitotoxicity or NMDA receptor overstimulation, leads to mitochondrial dysfunction and suppression of antioxidant enzymes, including GPX4^35,36^. Our findings suggest that glutamate excitotoxicity induced by NMDA receptor overactivation plays a pathological role in ApoE4-mediated cell damage. Additionally, lithium attenuates NMDA-induced ROS and lipid peroxidation and preserves GPX4 expression, pointing to a protective role against Ca^2+^-mediated ferroptosis initiation. Further, we demonstrated that lithium inhibits ApoE4-mediated InsP_3_R-1 upregulation. The robust effects of lithium to suppress InsP_3_R-1 upregulation is likely contributed by lithium inhibition of inositol monophosphatase, given that changes in levels of inositol trisphosphate or calcium influenced by inositol monophosphatase activity can affect InsP_3_R-1 expression^37^. This aligns with earlier reports that lithium can exert neuroprotective effects by modulating intracellular Ca^2+^ dynamics through inhibition of inositol monophosphatase and downregulation of NMDA receptor activity, in addition to blocking glycogen synthase kinase-3 activity to modulate gene transcription^15,19^. Furthermore, ApoE4 has been shown to increase Tau phosphorylation, potentially by activation of glycogen synthase kinase-3^38^. Lithium as an inhibitor of this kinase may decrease tau pathology in SAD expressing ApoE4/E4.

Mitochondrial respiration data revealed that lithium normalized aberrant increases in OCR and proton leak triggered by NMDA/RSL3 exposure in ApoE4/E4 cells, further supporting its role in maintaining mitochondrial bioenergetic homeostasis and reducing apoptosis-prone mitochondrial stress. In this context, lithium’s inhibition of oxidative stress and lipid peroxidation likely contributes to preserved mitochondrial function and overall cell survival. A schematic illustration of the effects of lithium on the pathological events in ApoE4/E4 iPSCs and potential underlying mechanisms is shown in Fig. 8.

**Figure 8.**
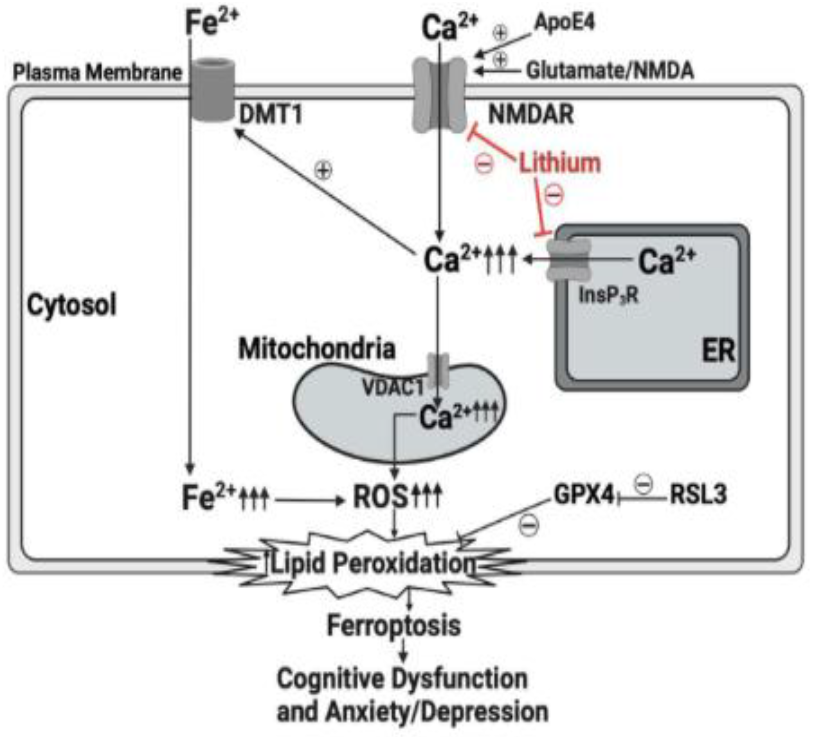
Lithium inhibits ApoE4/NMDA/RSL-3 mediated mitochondria dysfunction, oxidative stress and program cell death by ferroptosis. Excessive elevation of glutamate in Alzheimer’s disease (AD) activates N-methyl-D-aspartate receptors (NMDAR) pathologically, leading to excessive Ca^2+^ influx into cytosol via NMDARs. This can be aggravated by SAD risk factor ApoE4. The number and function of InsP_3_ receptors (InsP_3_R) on endoplasmic reticulum (ER) are pathologically increased in AD, leading to excessive Ca^2+^ release from the ER into the cytosol. Elevated [Ca^2+^]_c_ increases the expression of divalent metal transporter 1 (DMT1), an iron transporter, leading to the pathological elevation of cytosolic Fe^2+^ and further increases of ROS, lipid peroxidation and ferroptosis. RSL3 induces ferroptosis specifically by inhibiting glutathione peroxidase 4 (GPX4). Lithium provides robust protection against the upstream critical Ca^2+^ and Fe^2+^ dysregulation and associated mitochondria dysfunction, oxidative stress and program cell death by apoptosis or ferroptosis.

Our recent study also demonstrated that lithium inhibits pathological inflammation and programmed cell death by pyroptosis, likely by inhibiting upstream Ca^2+^ dysregulation in the brains of 5XFAD AD transgenic mice^10^. Together with the results in this study, lithium may provide neuroprotection against programmed neuronal death via its inhibition of both ferroptosis and pyroptosis in AD, although the molecular mechanisms, especially the role of Ca^2+^ dysregulation, warrant further investigation. It has been proposed that effective pharmacological treatment of AD should inhibit the critical upstream pathology or inhibit multiple pathologies, and lithium seems to fit these requirements. Although some studies have demonstrated the effectiveness of lithium at low clinical concentration (0.25-0.5 mM) in inhibiting mild cognitive function in AD patients^39^, more clinical studies are needed to confirm its effectiveness as either merely a symptom-relieving drug or as a disease-modifying drug for the prevention of AD.

One limitation of using lithium as a first-line drug to treat bipolar disease is its narrow therapeutic window^40^. The primary organ toxicity is its impairment of thyroid and kidney function^41,42^. Our recent study in mice demonstrated that intranasal lithium chloride in Ryanodex formulation vehicle (RFV)^43,44^ significantly increased lithium brain concentration, duration, and brain/blood lithium concentration ratio, compared to oral lithium chloride in RFV or intranasal lithium chloride dissolved in water10. Consistently, intranasal lithium chloride in RFV significantly inhibited memory loss and depressive behavior via inhibition of inflammatory pyroptosis in 5XFAD transgenic AD mice, without obvious adverse effects or organ toxicity towards the kidney or thyroid. In fact, intranasal lithium chloride in RFV protects kidney function in 5XFAD mice^10^. Together with the results demonstrating lithium’s protection against inflammatory pyroptosis in the current study, we propose that intranasal lithium in RFV may be an effective drug treatment for AD. As ApoE4 is a prominent risk factor for SAD, the capability of lithium to prevent mitochondrial and cell damage in ApoE4/E4 iPSCs from a SAD patient in this study suggests its potential to treat SAD patients, pending future clinical studies.

Overall, these results support a model in which lithium acts at multiple levels to prevent ferroptosis in ApoE4/E4 iPSCs from an SAD patient: (1) reducing iron uptake via DMT1 downregulation, (2) preserving lipid detoxification capacity through stabilization of GPX4, and (3) inhibiting Ca^2+^ and iron dysregulation associated oxidative stress, a key upstream trigger of ferroptotic death^37-39^. This multimodal action highlights lithium as a particularly promising therapeutic candidate for targeting ferroptosis in AD, especially in genetically susceptible populations with the ApoE4/E4 genotype, particularly in sporadic AD.

Despite these promising results, several limitations warrant discussion. The findings are derived from in vitro models, particularly in iPSCs, not neurons, and further validation in animal models or human brain tissue is required to establish translational relevance. Moreover, while the data support a role for Ca^2+^ in GPX4 regulation, the precise molecular intermediates involved, such as Ca^2+^-dependent transcription factors or signaling kinases, remain to be identified. Future studies should investigate whether lithium interferes with ferroptosis through direct modulation of GPX4 transcription, post-translational stabilization, or upstream signaling cascades.

## Conclusion

In conclusion, this study identifies lithium as a potent inhibitor of ferroptosis in human ApoE4/E4 iPSCs and provides mechanistic insight into its action via Ca^2+^-dependent regulation of GPX4, illustrating a molecular mechanism of ApoE4-mediated pathology in SAD patients. These findings suggest a neuroprotective mechanism for lithium and support its continued evaluation as a disease-modifying therapy in AD.

## Acknowledgements

This work was supported by grants to HW from the National Institute on Aging (R01AG061447) and NIA R01 Supplemental (3R01AG061447-03S1). The research was performed in the lab of Dr. Huafeng Wei and should be attributed to the Department of Anesthesiology, University of Pennsylvania. We appreciate technical support from Roland Zhang, an undergraduate student from the University of Pennsylvania and Justin Lu, a medical student from Drexel University, Philadelphia, U.S.A.

## Author Contributions

H.W. conceived and designed the study. Y.W., S.A., P.B., L.S., G.L. and H.W. conducted experiments and acquired the data. Y.W., S.A., P.B., L.S. and H.W. analyzed data. S.A., D-M.C and H.W. evaluated the experimental data and constructed the manuscript. All the authors reviewed and approved the final manuscript.

## Conflict of Interest

Huafeng Wei is an inventor for USA patents application by the University of Pennsylvania repurposing intranasal dantrolene nanoparticles to treat neurodegenerative diseases.

## Data Availability

Data is available upon request

## Notes

### Competing Interest Statement

The authors have declared no competing interest.

